# High contiguity long read assembly of *Brassica nigra* allows localization of active centromeres and provides insights into the ancestral *Brassica* genome

**DOI:** 10.1101/2020.02.03.932665

**Authors:** Sampath Perumal, Chu Shin Koh, Lingling Jin, Miles Buchwaldt, Erin Higgins, Chunfang Zheng, David Sankoff, Stephen J. Robinson, Sateesh Kagale, Zahra-Katy Navabi, Lily Tang, Kyla N. Horner, Zhesi He, Ian Bancroft, Boulos Chalhoub, Andrew G Sharpe, Isobel AP Parkin

## Abstract

High-quality nanopore genome assemblies were generated for two *Brassica nigra* genotypes (Ni100 and CN115125); a member of the agronomically important *Brassica* species. The N50 contig length for the two assemblies were 17.1 Mb (58 contigs) and 0.29 Mb (963 contigs), respectively, reflecting recent improvements in the technology. Comparison with a *de novo* short read assembly for Ni100 corroborated genome integrity and quantified sequence related error rates (0.002%). The contiguity and coverage allowed unprecedented access to low complexity regions of the genome. Pericentromeric regions and coincidence of hypo-methylation enabled localization of active centromeres and identified a novel centromere-associated ALE class I element which appears to have proliferated through relatively recent nested transposition events (<1 million years ago). Computational abstraction was used to define a post-triplication *Brassica* specific ancestral genome and to calculate the extensive rearrangements that define the genomic distance separating *B. nigra* from its diploid relatives.

---

Decoding complete genome information is vital for understanding genome structure, providing a full complement of both the genic and repeat repertoire and uncovering structural variation. Such information also provides a foundational tool for crop improvement to facilitate the rapid selection of agronomically important traits and to exploit modern breeding tools such as genome editing^1,2,3^. Whole genome duplication and abundant repeat expansion has led to an approximate 660-fold variation in genome size among angiosperms^4^ and in particular the low complexity of repetitive regions create challenges for complete genome assembly using short read sequence data^5^.

Recently advances in long read sequencing technologies such as Pacific Biosciences (PacBio) and Oxford Nanopore Technology (ONT)^6^ which combined with genome scaffolding methods such as optical mapping and chromosome conformation capture (Hi-C) have led to a paradigm shift in our ability to obtain complete and contiguous genome assemblies^7,8,9^. Both approaches can produce remarkably long reads; although, the error rate is significantly higher than more accurate Illumina short reads, which until recently limited their use to scaffolding to improve assembly contiguity^10^. However, correction algorithms to reduce error rates, and recent technological improvements have increased the output and quality of ONT sequence data making routine and cost effective assembly of large eukaryotic genomes possible^11^.

The *Brassicaceae* is an important plant family with approximately 3,800 species including commercially important vegetable, fodder, oilseed and ornamental crops. The *Brassiceae* tribe has a history of extensive whole genome duplication events, including the *Brassica* genus specific whole genome triplication (WGT), which occurred approximately 22.5 million years ago (Mya)^12,13^ and is assumed to be shared by the three important diploids (*B. rapa*, AA, 2n=2x=20; *B. nigra*, BB, 2n=2x=16; and *B. oleracea*, CC, 2n=2x=18) that form the vertices of U’s triangle^14^. Among these three, *B. nigra* (B-genome) has been neglected with regards to both genetic analyses and selection through breeding. Due to its limited domestication and its production as out-crossing populations it has retained valuable allelic diversity compared to its relatives, making it an untapped repository for *Brassica* breeding^15^. Among the six species of U’s triangle, five have been sequenced including most recently *B. nigra*, but the assemblies cover at most 80% of the estimated genome size and almost all were very highly fragmented due to the use of short-reads alone^16,17,18,19,20^. Recently, the *B. rapa* reference genome was improved using PacBio sequencing^19^ and one genotype each of *B. rapa* and *B. oleracea* was sequenced using a combination of ONT and optical maps demonstrating the utility of these technologies for complex duplicated genomes^21^.

The work described represents the nearly complete assembly of two *B. nigra* genomes (Ni100 and CN115125) using a combination of ONT sequencing, Hi-C and genetic map-based scaffolding. A short-read assembly of Ni100 allowed comprehensive benchmarking of the long-read assemblies. Remarkably, direct methylome profiling utilising the ONT data allowed candidate active centromeres of the chromosomes to be resolved, a feature previously unannotated in short read assemblies. In addition, computationally defined genomic distances between the three *Brassica* diploid genomes allowed the construction of an ancestral *Brassica* specific genome.

## Results

A combination of nanopore sequencing, Illumina error correction, chromosome confirmation capture (Hi-C) and genetic mapping was used to generate two *de novo* assemblies for the diploid *Brassica* species, *B. nigra* (Ni100 and CN115125 genotypes). Identical sequential steps were followed to assemble the contigs for each genome including the development of high-quality sequencing data sets, genome assembly and polishing with short reads (Supplementary Figure 1). After testing a number of published assembly software pipelines (Supplementary Table 1) the final contigs were derived from SMARTdenovo using 30-64x coverage of CANU^22^ corrected reads.

Although largely context dependent, nanopore sequence data can show error rates up to 15%. Thus, sequence correction was completed using eight rounds of Pilon^23^ with approximately 100x coverage of Illumina data, quality was assessed at each round through Benchmarking Universal Single Copy Orthologue (BUSCO)^24^ scores and qualimap^25^ (Supplementary Figure 2; Supplementary Table 2). For both genotypes the read alignment rate was high (>98%) and both tools indicated a significant improvement after correction suggesting final error rates of 0.8 % (CN115125) to 0.2% (Ni100) at the base pair level. The two long read (LR) assemblies were generated over a period of approximately 12 months, during which time ONT upgraded their library construction kits, pore chemistry and base calling software. The combined impact of which was noted in an overall improvement in quality, average read length and usable data output for Ni100 and in final assembly contiguity (Supplementary Table 3, 4 and 5). The CN115125 assembly was more fragmented (cf. a contig N50 length of 0.288 Mb and 17.1 Mb) thus scaffolding using proximity ligation, a combination of Chicago^™^ and Hi-C, was used to improve the contiguity up to 193 fold, with a final N50 length of 55.7 Mb (Supplementary Figure 3). In both instances genetic anchoring was used to generate the final chromosome-scale assemblies of the two *B. nigra* genotypes, CN115125 (C2-LR) and Ni100 (Ni100-LR) (Table 1, Figure 1, Supplementary Figure 4, Supplementary Table 6).

**Figure 1.**
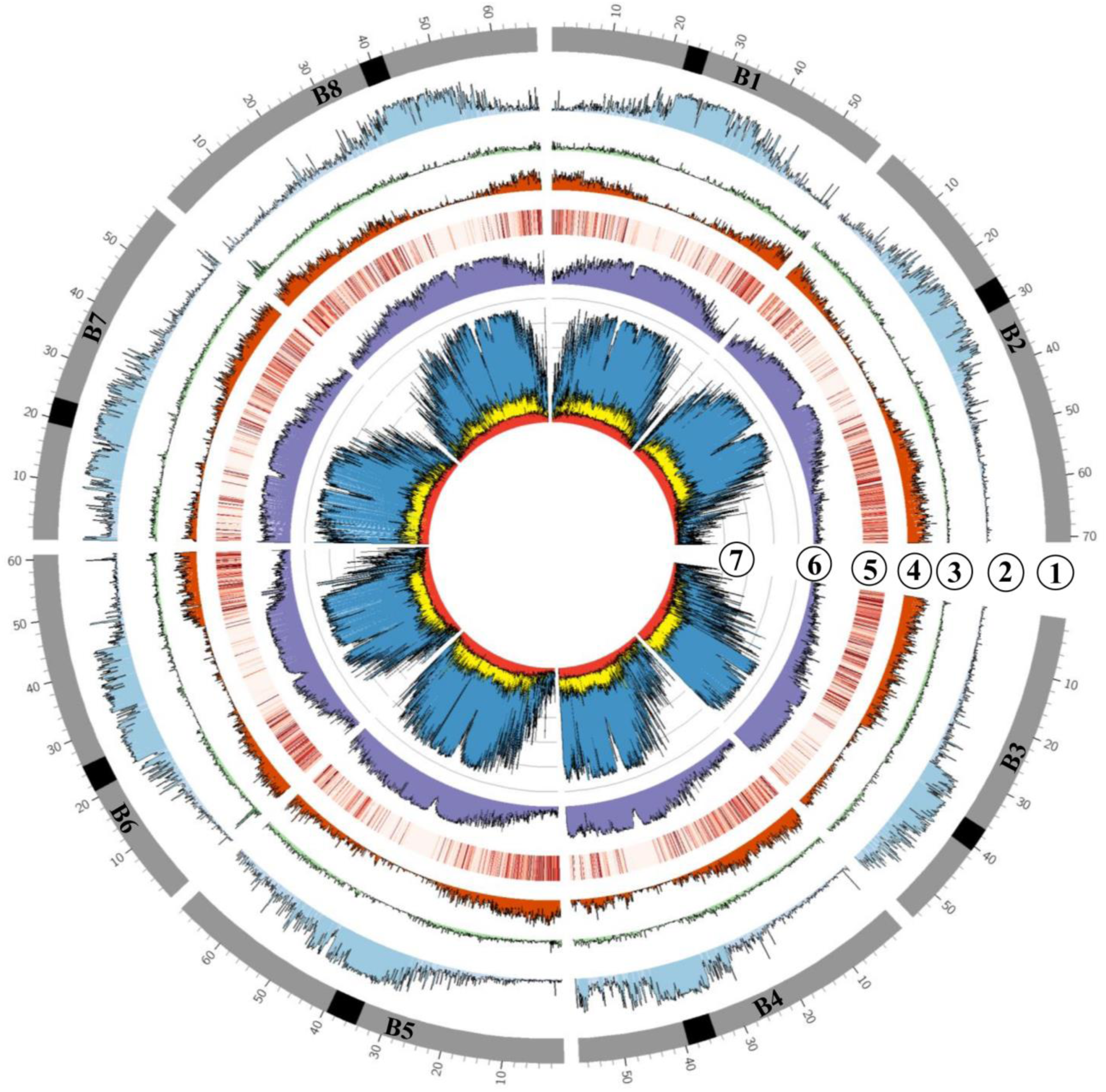
Genomic features of the *B. nigra* Ni100-LR assembly. Tracks: (1) Chromosomes with centromere (black band); (2) Class I retrotransposons (nucleotides per 100 Kb bins); (3) Class II DNA repeats (nucleotides per 100 Kb bins); (4) Gene density (genes per 100 Kb bins); (5) Gene expression in leaf tissue (log^10^(average TPM) in 100 Kb bins); (6) ONT CG-methylation profile (ratio per 100 Kb); (7) Whole genome bisulfite methylation profile (nucleotides per 100 Kb bins), CG-blue, CHG-yellow, CHH-red.

**Table 1.** Statistics of the *B. nigra* genome short-read (SR) and long-read (LR) assemblies.

| **Assembly** | **Ni100-SR** | **Ni100-LR** | **C2-LR** |
| --- | --- | --- | --- |
| Estimated genome size (k = 17), Mb | 570 | 570 | 608 |
| Assembly size, Mb | 447 | 506 | 537 |
| No. of chromosomes | 8 | 8 | 8 |
| Genome coverage | 78% | 89% | 88% |
| Number of sequences | 19,203 | 58 | 963 |
| Longest scaffold, Mb | 53 | 50 | 71 |
| Scaffold N50, Mb | 43.9 | 60.8 | 55.7 |
| Scaffold L50 | 5 | 4 | 5 |
| Contig N50, Kb | 48 | 17,127 | 288 |
| Ambiguous bases “N”, Kb | 33,737 | 13 | 390 |
| BUSCO: % Complete | 97 | 97 | 94.4 |
| % GC content | 37 | 37 | 37 |
| Number of genes<sup>1</sup> | 56,331 | 59,877 | 67,030 |
| *High-confidence genes (%) | 55,141 (98) | 57,798 (97) | 64,071 (96) |
| Low-confidence genes (%) | 1,190 (2) | 2,079 (3) | 2,959 (4) |
| Repeats and TE space, Mb (%) | 183 (41.5) | 273 (54) | 263 (49) |
| Uncharacterized genome, Mb (%) | 125.5 (22) | 64 (11) | 71.4 (12) |

A short read Illumina *de novo* assembly for *B. nigra* genotype Ni100 (Ni100–SR) was used for further validation of the nanopore assemblies. The Ni100-SR assembly has a total length of 446.5 Mb from 19,195 contigs, of which 367.5 Mb was anchored to eight pseudo-chromosomes (Table 1; Supplementary Table 6). Alignment and visualisation of corresponding pseudo-chromosome sequence from the three *B. nigra* assemblies revealed high levels of collinearity (Figure 2A). A number of inversions were noted, in particular a large inversion at the bottom of B4 distinguished the SR assembly. The B4 region was difficult to scaffold in the SR assembly due to limited recombination and was largely ordered exploiting synteny data from *Arabidopsis thaliana*. It was apparent that there was expansion of the ONT assemblies in regions presumed to be pericentromeric, as shown in Figure 2A and 2B. The level of coverage of these regions also varied between the long read assemblies.

**Figure 2.**
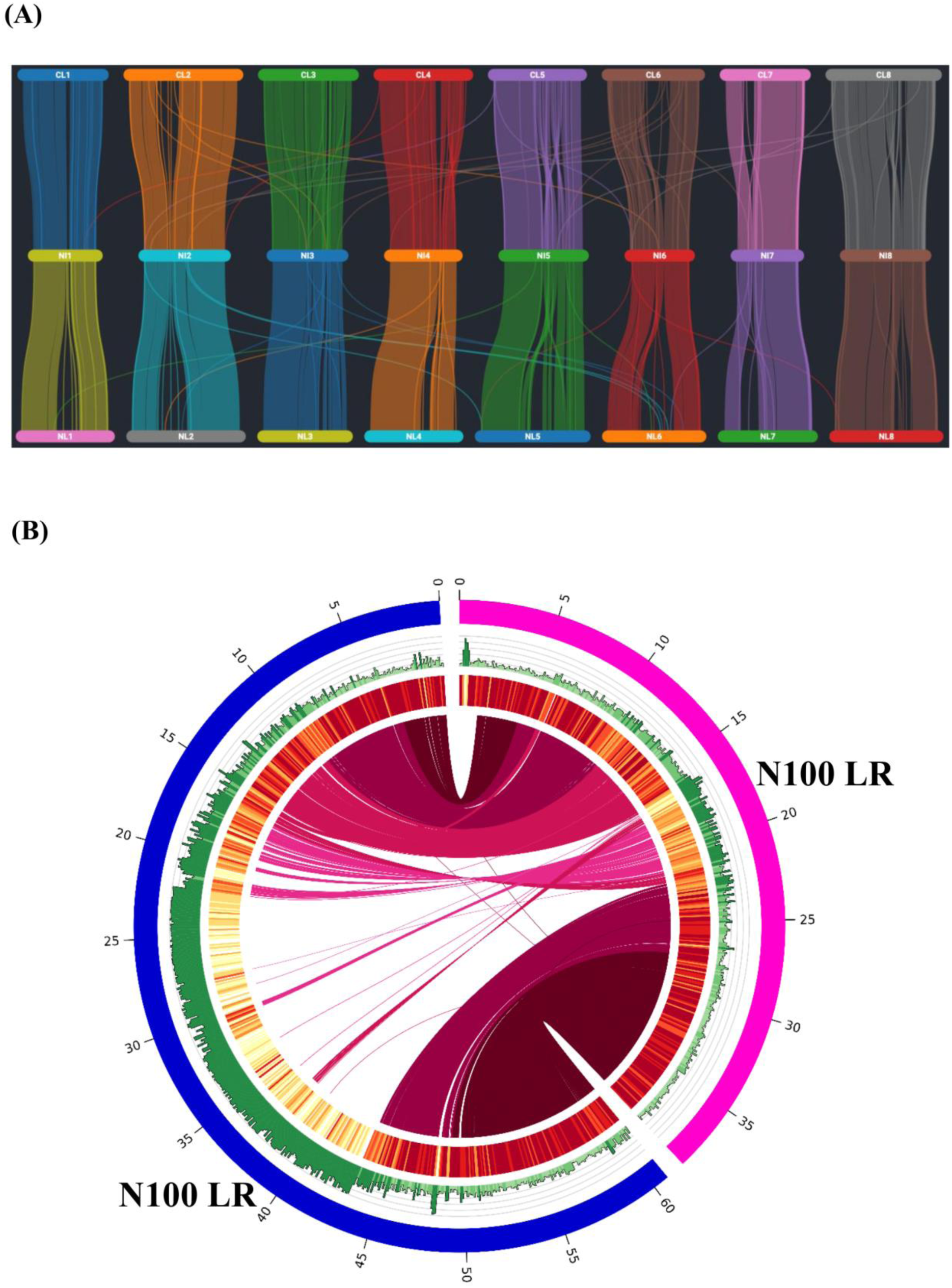
Comparison of the *B. nigra* assemblies. (A) Chromosome level genome alignment of the Ni100-SR (NS) assembly (centre) against the long read assemblies, CN115125 (top) and Ni100 (bottom). The plot was created using Synvisio (https://github.com/kiranbandi/synvisio). (B) Alignment of the short (NS) and long-read (NL) assemblies for chromosome B5 of Ni100.

Gene annotation from the two LR and two SR assemblies (Ni100-SR and the previously published YZ12151^20^) were rationalized to generate a final *B. nigra* gene complement of 67,030 and 59,877 gene models in the two genotypes, CN115125 and Ni100, respectively. These numbers are in line with the predicted pan-gene content of diploid *B. oleracea*, with 63,865±31 genes^26^. An additional 3,546 genes were annotated in Ni100-LR compared to the Ni100-SR assembly. A homology search performed using GMAP^27^ (minimum identity and coverage of 95%) indicated only 914 of the additional genes were unique to the Ni100-LR assembly (Supplementary Figure 5A). This discrepancy was due to both co-assembly of highly similar genes in the SR data and assembly errors that precluded accurate gene annotation. Read mapping of Illumina data back to the SR and LR assemblies showed a marked increase of 9% multi-mapping reads in the latter, with a concomitant reduction in non-concordant matches, suggesting the resolution of duplicated or highly homologous sequences in the LR assembly (Supplementary Table 7). The recent ONT assemblies of *B. rapa* and *B. oleracea* studied the self-incompatibility locus, or the S locus region, which due to its repetitive structure has been notoriously difficult to assemble, to infer the enhanced contiguity of the LR derived genome sequences^28^. The S locus region was identified and compared in the two *B. nigra* long read assemblies showing complete assembly of two differing S locus haplotypes (Supplementary Figure 6). A comparison between the two ONT assemblies would be expected to identify such genotype differences. Along with approximately 10% of the annotated genes being specific to either assembly, the CN115125 genotype showed a higher prevalence of tandemly and proximally duplicated genes (Supplementary Figure 5C and Supplementary Table 8 and 9).

In order to investigate the global gene content across the genomes, annotated genes from the three representations of *B. nigra* were clustered with genes from *B. rapa, B. oleracea* and *A. thaliana* using Orthofinder^29^ (Supplementary Figure 5D). From 311,510 genes included in the cluster analysis, 293,144 genes were grouped into 46,271 gene clusters, of which 17,464 clusters contained genes from all six genomes (Supplementary Figure 7A; Supplementary Table 10). The diploid *Brassica* species ranged in number of species-specific genes with *B. rapa* containing the least while *B. nigra* contained the most (Supplementary Figure 7B,C; Supplementary Table 10). In *B. nigra* 936 and 603 gene families appeared to be expanded or contracted, respectively compared to their diploid relatives (Supplementary Figure 8A). A similar pattern of expansion (970) and contraction (547) was also observed for the *B. rapa/B. oleracea* lineage. Sixty-nine of the *B. nigra* specific gene families were deemed to be rapidly evolving by CAFÉ^30^ and functional analyses demonstrated the genes were enriched in response to abiotic or biotic responses, structural molecule activity and unknown molecular functions (Supplementary Figure 8B). Since it is often noted that families related to stress are more prone to differential copy number variation, differences in R-genes, transcription factors (TF) and protein kinase families were assessed in each of the genomes. The distribution of R gene families across the species appeared to be directly related to genome size and/or expansion of the transposable element complement, with *B. oleracea* showing the largest expansion of R genes. CN115125 in particular showed increased membership of TF families, with both *B. nigra* genotypes showing a higher prevalence of B3, C2H2 and NAC domain TFs compared to their diploid relatives (Supplementary Table 11, 12 and 13).

Structural variations (SVs) including deletions, insertions, duplications, inversions, and translocations that differentiate genotypes were cataloged between both genomes using ONT-reads. The raw ONT reads from Ni100 and CN115125 were aligned to both LR assemblies and SVs were quantified using two different SV callers (Sniffles^31^ and Picky^32^). Self-alignment was used to estimate a false positive rate for each genome, which was higher for the CN115125 assembly (cf. 6,307 to 2,230 events) (Supplementary Table 14; Figure 3). High quality SVs were considered to be those identified with both software packages (Figure 3D and 3E). At least fifteen random SVs of each type were assayed manually and suggested almost 100% prediction accuracy for deletions and insertions; however, some of the larger predicted events seemed less reliable and it was apparent that a number of deletions were overlooked suggesting the criteria may have been too stringent. Overall, 7,181 SV were identified by comparing the CN115125 ONT reads against the Ni100-LR reference (C2onNL) (Supplementary Table 14; Figure 3). Among the 7,181 SVs, 70% (5,059) were deletions, of which 45% were closely associated with genic regions (Supplementary Table 14; Figure 3B). Likewise, 6,078 SVs were found by mapping Ni100 ONT reads against the C2-LR assembly (NLonC2), with deletions (3,856) again prevailing, followed by insertions (1,921) (Figure 3A). In general deletions were more prevalent, with shorter deletions and insertions (< 1Kb) being more evenly balanced. Since the reciprocal read mapping should identify effectively the same events this could suggest limitations to the automated pipeline for curating larger rearrangements, but this would be exacerbated by an overall average read length in the CN115125 ONT reads (10.9 Kb cv 20.4 Kb) and some variation in genomic regions assembled (Supplementary Table 4).

**Figure 3.**
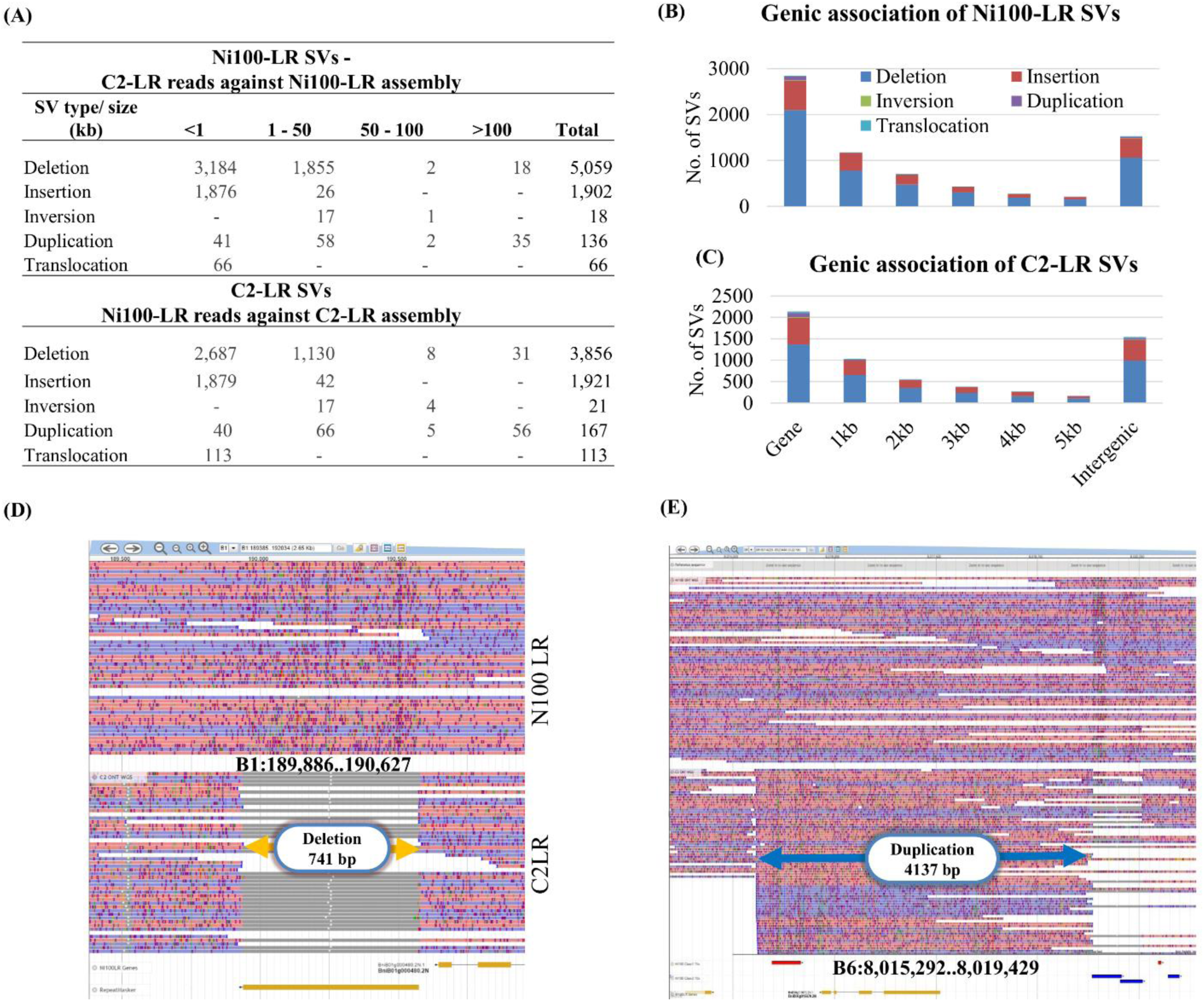
Identification of Structural variants (SVs) in *B. nigra* Ni100-LR and C2-LR. (A) Summary of consensus SVs (identified by Sniffles and Picky). Association of Ni100-LR SVs (B) and C2-LR SVs (C) with gene regions and flanking DNA. Jbrowse alignments showing a deletion (741 bp) of a transposable element (D) and duplication (4137 bp) (E) of an annotated gene in the C2-LR genome compared to Ni100.

A *Brassica* B genome specific repeat library with 1324 families was developed using multiple annotation tools and was used to survey the repetitive genome fraction of the long-read (Ni100-LR, C2-LR) and short-read *B. nigra* assemblies (Ni100-SR and YZ12151-SR) (Supplementary Table 15). Repeats spanned 49% and 54% of the CN115125 and Ni100-LR genome assemblies, respectively, compared to 33% (YZ12151) and 41% (Ni100) in the two short read assemblies. The increase in repeat content of the long read assemblies predominantly resulted from a rise in annotated Class I transposons, in particular Gypsy and Copia elements, which increased by 27.8% and 40.5%, respectively in the Ni100 assembly (Supplementary Table 15). The distribution of repeats revealed class I transposons were more common in traditionally heterochromatic regions such as centromeric, pericentromeric and sub-telomeric regions, while class II DNA transposons were more evenly distributed across the genome (Figure 1; Supplementary Figure 4). The identification of centromere and telomere specific repeats suggested the ONT assemblies provided more complete access to the chromosome structure (Supplementary Figure 9). The repeat fraction appears to reflect the estimated genome sizes of the diploid *Brassicas* with *B. nigra* lying between *B. oleracea* (~60%)^21^ and *B. rapa* (~38%)^19,21^.

Almost all families were similarly distributed in the two long-read assemblies apart from LTR-Gypsy elements, which were ~5% higher in Ni100, suggesting either Ni100 specific amplification or assembly of these elements (Supplementary Table 15). Full-length long terminal repeat retrotransposons (FL-LTR-RTs) were annotated and compared in Ni100-SR and the two long-read assemblies. A total of 1220, 4491 and 3381 FL-LTR-RTs were identified in Ni100-SR, Ni100-LR and C2-LR assemblies, respectively, with an average length of ~6 Kb (Supplementary Table 16; Supplementary Figure 10A). The increased annotation of such elements in the long read assemblies indicates the benefits of the technology for assembling low complexity redundant sequences. Based on repeat domain protein homology the FL-LTR-RTs were grouped into 14 different families, where 41-44% had homology with known Gypsy families, 38-42% with Copia families, and 13-20% were unknown FL-LTR-RTs (Figure 4A). Notably, among the 14 FL-LTR-RTs families, members of the ALE (Copia) and OTA (Gypsy) families were specifically increased in copy number in the long read assemblies and more so in the Ni100-LR assembly (Figure 4A; Supplementary Table 16; Supplementary Figure 10B).

**Figure 4.**
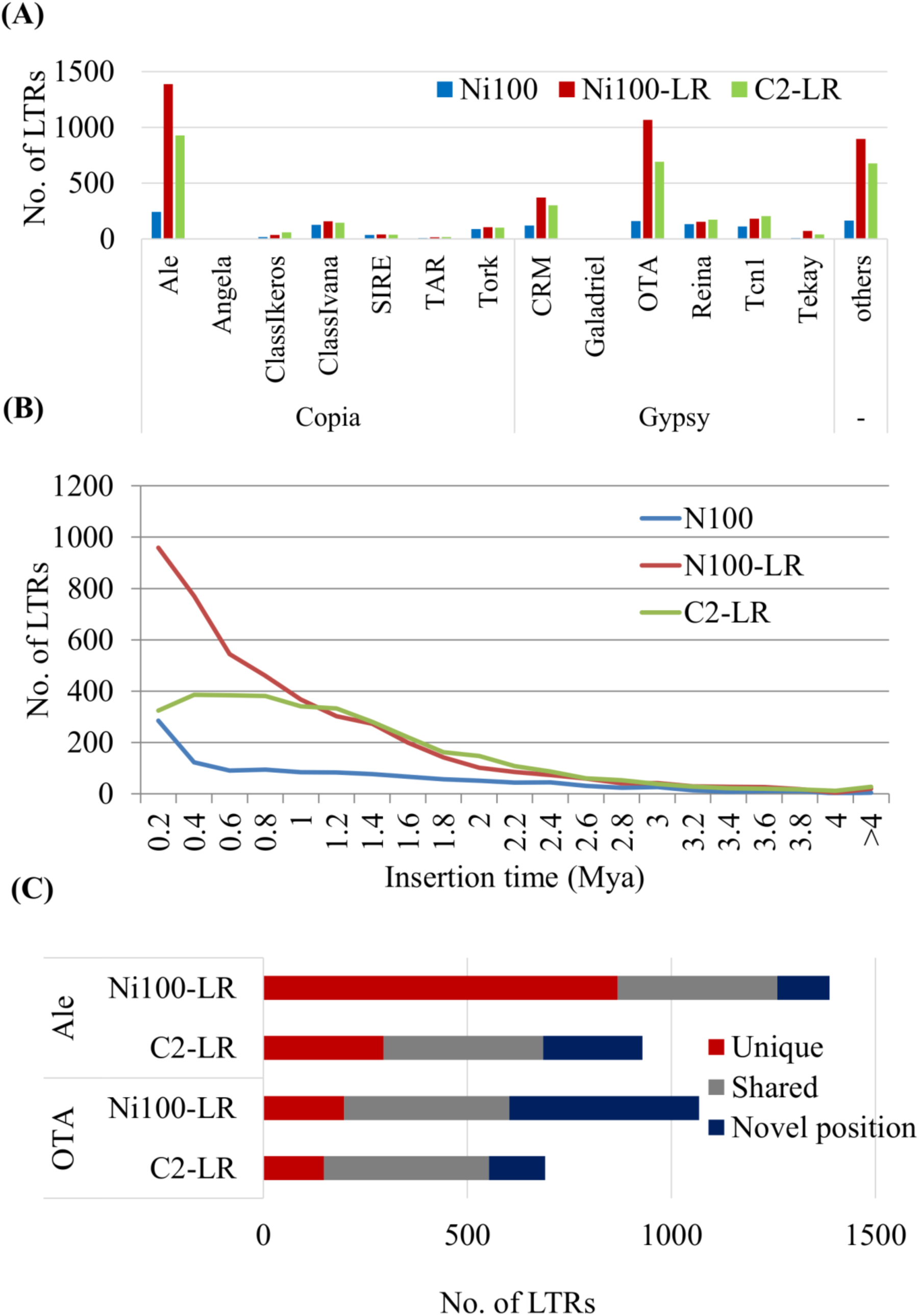
Annotation of full-length long terminal repeat retrotransposons (LTR-RT) in *B. nigra* genomes. (A) Copy number of FL-LTR-RTs from 14 different families. (B) Age distribution of FL-LTR-RTs in three *B. nigra* assemblies. (C) Comparison of insertion sites of two FL-LTR-RTs (Ale and OTA) in the ONT assemblies.

The age distribution analysis of FL-LTR-RTs elements based on divergence of the LTR region showed recent amplification of LTRs in both genomes. About 91% (4068) and 86% (2912) of the FL-LTR-RTs in Ni100-LR and C2-LR assemblies, respectively were amplified less than 2 Mya (Figure 4B; Supplementary Figure 11; Supplementary Table 16), with more recent and continuous proliferation of LTRs (3056, 68%) aged less than 1 Mya in Ni100-LR compared to C2-LR. ALE family elements showed more recent amplification (< 0.2 Mya) in Ni100-LR compared to C2-LR, while OTA LTRs showed a more consistent pattern between the two genomes. Analysis of the insertion sites revealed 405 OTA (59%) and 391 ALE (42%) were conserved in both genomes, and in line with the increase in density there were a higher percentage of uniquely inserted ALE elements in Ni100-LR genome compared to C2-LR genome (Figure 4C). A phylogenetic analysis of the ALE and OTA FL-LTRs suggested Ni100 genotype specific amplification of particular members of each family (Supplementary Figure 12).

ONT technology allows direct identification of base modifications such as 5-methylcytosine (5-mC)^33^, although this had yet to be shown in plant genomes. Nanopolish was used to detect 5-mC in the CG context in the Ni100 ONT unassembled reads, which provided higher coverage and quality compared to the CN115125 reads. The resultant calls were compared with methylation status detected using whole-genome bisulfite sequence data and showed excellent correlation irrespective of filtering for quality of call (R-values range from 0.93 – 0.97) (Figure 5B-D). As perhaps expected, the observed methylation showed similar patterns to those detected for related *Brassica* diploids^16^; with a higher prevalence of 5-mC in repeat sequences and lower methylation rates across annotated gene bodies (Figure 1; Supplementary Figure 13). Of note, there were islands of reduced methylation observed for each chromosome that were also associated with lower gene and higher repeat density regions. Centromeric regions have been associated in *Brassica* species and more specifically in *B. nigra* with particular sequences including CRB (centromeric retrotransposon of *Brassica*) and a B-genome specific short repeat fragment (pBN 35)^34,35^. The distribution of these centromere-associated repeats aligned with the detected hypomethylated regions. Furthermore, members of the more prevalent ALE family, which also has >70% homology with CRB, localized to the same region (Figure 5A). More recently sequences identified through interaction with the centromere specific histone CENH3, have been sequenced for *B. nigra*, which co-aligned with the hypomethylated regions, suggesting capture of much of the active centromere^36^.

**Figure 5.**
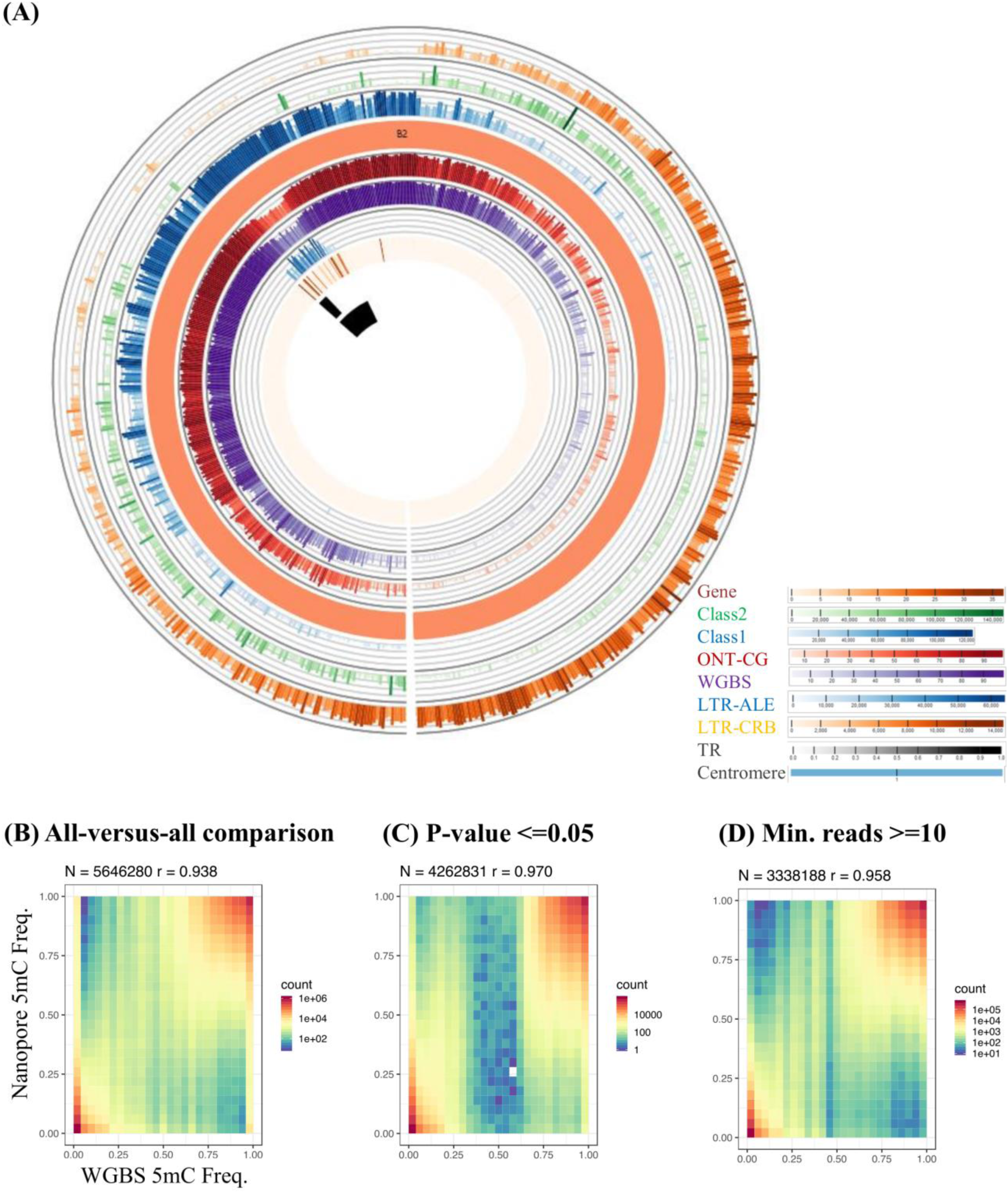
Comparison of methylation data from whole genome bisulfite (WGBS) and ONT sequencing in Ni100. (A) Genome features of the B2 chromosome of the Ni100-LR assembly, from outer to inner circle: gene density; Class II retrotransposon; Class I DNA transposon; chromosome cartoon; methylation profile from ONT data; methylation profile based on WGBS; ALE copia; centromeric retrotransposon of *Brassica* (CRB); *B. nigra* specific centromeric tandem repeat; putative centromere region. (B-D) Comparison of 5-methyl cytosine frequency detected by WGBS and ONT; frequency distribution plot without filtering (B) and filtering based on calls with a p-value ≤ 0.05 (C) or minimum ONT read depth of 10 (D).

Although the analyses of nested LTRs has generally been limited to cereal genomes, they would be expected to play a significant role in evolution of chromosome structure and specifically repeat dense regions such as centromeres. ALE LTRs were prominent in the centromeric regions and showed high levels of nested insertion (Figure 6). Overall 262 nested TE events were found throughout the Ni100-LR genome of which 68% (179) were in centromeric regions. Across all events, most involved two LTRs while 10 events involved >2 LTRs (Figure 6B; Supplementary Table 17). In-depth characterization of nested TEs in the centromeric region of chromosome B5 revealed 24 out of 26 of the nested TE events were created by ALE-LTRs, and all bar one of the events involved the same family member inserting into the host LTR. The predominantly young age (< 1 Mya) of the nested elements suggests continuous and recent rearrangement of the centromeric regions by this mechanism (Figure 6B).

**Figure 6.**
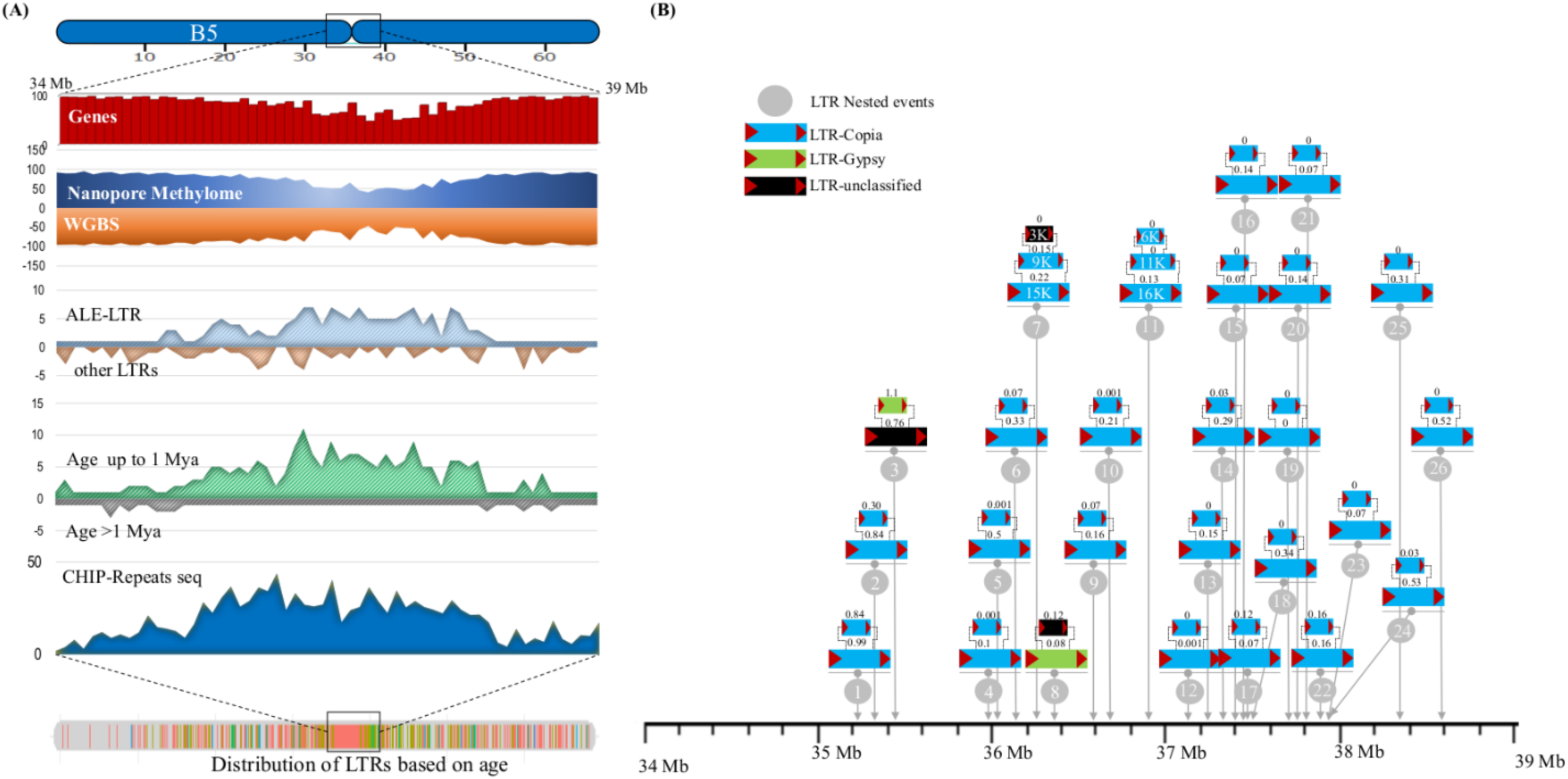
Characterization of centromeric region of chromosome B5 of Ni100-LR genome. (A) Distribution of various genomic features on the 5 Mb centromere region, including genes, methylome (ONT and WGBS), full-length LTRs (ALE LTR and other 13 family LTRs), distribution of young (<1 Mya) and old LTRs (>1 Mya), distribution of CENH3 repeats. (B) Nested insertion of full-length LTRs in the centromeric region. Age in Mya is shown above each element.

Much effort has been placed on defining an ancestral Crucifer genome that predates the supposed *Brassica* specific WGT event ^37^. Ancestral karyotype blocks were constructed for C2-LR and Ni100-LR based on shared gene content and order for orthologous copies of each *A. thaliana* gene (Supplementary Figure 14 and Supplementary Table 8). Based on the two-step mode of genome evolution inferred from the genomes of *B. rapa* and *B. oleracea*^38^, which is predicated on genome dominance in newly formed polyploids, as expected the blocks were found predominantly in three copies but with biased genic content. The least fractionated genome maintains approximately 70% of the orthologous gene copies, while the most fractionated 1 (MF1) and MF2 retain approximately 49% and 42%, respectively (Supplementary Figure 5C). A phylogenetic analysis of the triplicated orthologues confirmed a shared WGT among the *Brassica*s with genes from across the three species of each triplicated genome being more similar than those within the same species (Figure 7C). Some smaller genomic regions were found in additional syntenic blocks in each genome, which could represent more ancient whole genome duplication events or further localized segmental translocations. These supplementary blocks were more prevalent in the CN115125 genome and could explain the higher prevalence of duplicated genes in this genome (Supplementary Figure 5C; Supplementary Table 8).

**Figure 7.**
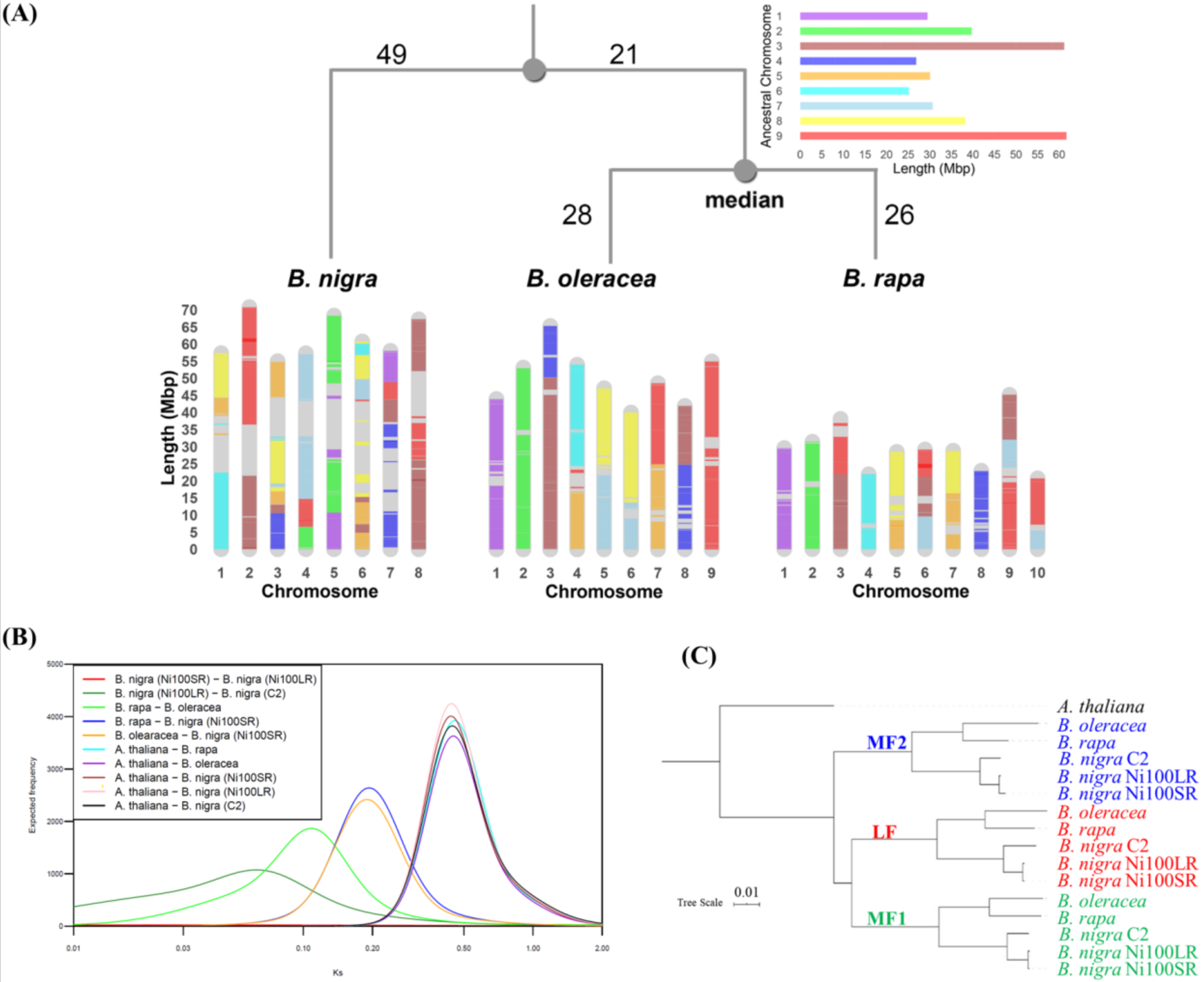
Genome rearrangements and evolution of *Brassica* species. (A) Development of *B. rapa, B. nigra* and *B. oleracea* genomes based on ancestral genome. Blocks are “painted” with colours corresponding to the ancestral chromosomes. (B) Divergence time estimation based on Ks (synonymous substitutions) distributions. Gaussian mixture models fitted to frequency distributions of Ks values obtained by comparing pairs of syntelogs between different *Brassica* species or the subgenomes of each species are shown. (C) Phylogenetic relationship between the subgenomes of different *Brassica* species. A maximum likelihood tree constructed based on concatenated sequences of 1150 syntelogs between *A. thaliana* and each of the subgenomes (LF, MF1 and MF2) of the *B. rapa*, *B. oleracea* and *B. nigra* is presented. Clade support values near nodes represent bootstrap proportions in percentages. All unmarked nodes have absolute support.

Genomic differences or similarities among species, as well as the mechanisms by which genomes evolve, can be identified by comparing the order in which genes or syntenic blocks appear in both close and distant relatives ^39^. The changes to block orders are defined in terms of certain rearrangement operations both within a chromosome or between chromosomes, such as reversal, transposition, fusion, fission and translocations. These types of operations, which occur often throughout evolution of a species can be abstracted computationally as a series that results in a change to the linear ordering of genes. One approach to measure the degree of dissimilarity between species is to find the series with the fewest possible operations, the most parsimonious evolutionary process or the “genomic distance”, that transforms the genome from one “version” to another. In the Double-Cut-and-Join (DCJ) model ^40^, two genomes being compared are represented as a “breakpoint” graph, allowing the metric the DCJ distance between genomes to be calculated. The pairwise DCJ distances between the three *Brassica* diploid genomes were thus calculated to be: *d_rapa,nigra_* = 96; *d_nigra,oleracea_* = 98; *d_rapa,oleracea_* = 52. In addition to measuring genomic difference or similarity, the order of blocks in extant genomes provides rich information that can be used in reconstruction of ancestral gene orders. The median problem for genome rearrangements is the simplest instance of a problem of reconstructing the gene order of an ancestral species. Given gene orders in a set of genomes G and a distance measure d, the median problem finds a genome m that minimizes the sum of distances *d^Σ^*= Σ_g∈G_ *d(m, g)*. The ASMedian-Linear algorithm ^41^ was used to calculate the median *Brassica* genome as nine ancestral chromosomes with a genome size of 321 Mb (Supplementary Figure 15; Supplementary Table 18) consisting of 178 blocks. Each block in the median genome was mapped to the three extant genomes as shown in Figure 7A, while Supplementary Figure 16 shows the detailed position of each ancestral median block and their relative orientation. The calculated median genome *m* under the DCJ model generates a minimum total DCJ distance: d^Σ^= Σ_g∈G_ d_m,g_= 124, where *d_m,rapa_* = *26 d_m,nigra_* = *70*, and *d_m,oleracea_* = 28. Given these distances, a rooted ultrametric phylogenetic tree was approximated (Figure 7A), where the position of the median minimizes the total DCJ distance *d^Σ^*. Based on molecular clock theory it can be seen that the median is the most recent common ancestor of *B. rapa* and *B. oleracea*, while the overall ancestor is almost 1/3 of the way along a path from the median to *B. nigra*. The DCJ distance between the genomes corresponded with the age of divergence estimated from the synonymous substitutions (Ks) rates among the coding regions of orthologous gene pairs across the genomes, with *B. oleracea*/*B. rapa* having diverged from *B. nigra* some 11.5 Mya, while they diverged from each other only 6.8 Mya (Figure 7B; Supplementary Table 19).

## Discussion

Recent advancements and cost reductions in long-read sequencing technologies are facilitating the generation of high-quality genome assemblies even for species that have evolved through recursive whole genome duplication events (WGD)^42^. High quality and highly contiguous assemblies were generated for two genotypes of the mesopolyploid *B. nigra* using nanopore sequencing, chromosome level scaffolding with Hi-C and genetic mapping data. Remarkably, the final contig N50 length was 17.1 Mb (Ni100-LR), one of the longest among the 324 plant genomes published to date (Supplementary Figure 17; Supplementary Table 20). Comparing the two ONT assemblies, the Ni100-LR assembly was better in terms of contiguity and capture of repeat rich centromeric regions, reflecting rapid improvements in the technology and suggesting the importance of both read length (11 kb cv 20 kb) and read coverage (29x cv 64x). Accurate quantification of errors in the Ni100 nanopore assembly through comparison with an Illumina short-read assembly of the same genotype suggested an accuracy of 99.986%, which was only improved marginally (99.998%) with eight rounds of short-read polishing suggesting that nanopore reads can provide highly accurate assemblies of complex genomes. The error rate was higher for the CN115125 assembly (0.8% cv 0.2%), again reflecting the recent improvements in ONT technology.

The recognition that both small (copy number, presence/absence) and large (chromosomal rearrangements) structural variation (SV) plays an important role in controlling key agronomic traits is gaining traction^43^; yet deciphering such variation with short-read data has proved problematic^44^. Long-read sequencing technologies have distinct advantages in predicting SVs^45^, however the current limitation is developing and training software to accurately identify such variants. The current analyses utilized two widely accepted software tools for detecting SVs and cross-validation was attempted to improve the accuracy of the calls. The large difference in the number of events discovered by the different protocols probably reflects a combination of a higher false and a lower positive discovery rate between the two. Considering only the cross-validated calls, a large number of events differentiated the two *B. nigra* genotypes, and many would impact gene expression and potentially phenotype, thus underlying the need for improved tools for SV analyses to capture this valuable information.

It is well established that long-read sequence data provides a more comprehensive coverage of the genome^46^, perhaps most obviously reflected in the increased capture of low complexity repeat sequences. Repeat analysis revealed about 14% more repeats in the long read assembly of Ni100 compared to the short read assembly (54% vs 41.2%) and in particular a more complete assembly of the repeat rich centromeric and peri-centromeric space. Centromeres are essential structures for the maintenance of karyotype integrity during meiosis, ensuring fertility of developed gametes through strict inheritance of full chromosome complements; yet centromeres still remain underexplored, especially in larger genomes. Although the active centromere is incredibly diverse in size and sequence among species, it is characterized through its cohesion with the centromere-specific histone H3-like protein, CENH3, and it has been suggested that association with CENH3 is controlled through epigenetic means, including a decrease in CG methylation^34^. Direct CG methylation profiling utilizing the ONT data suggested not only the efficacy of this approach (93-97% correlation with WGBS) but also demarcated the active centromere in the assembly, with hypo-methylated regions being co-located with known and novel centromeric repeat sequences. At least three of the chromosomes for Ni100 (B1, B3 and B8) showed multiple hypomethylated islands within or adjacent to the putative centromere region, which also coincided with centromeric specific repeats (Supplementary Figure 18). It was noted for *B. rapa* that such repeats found outside the presumed centromeric region may be evidence of ancient paleo-centromeres, remnants of the WGD events^19^. However, all additional sites coincided with hypo-methylation suggesting functionality of the regions. This could imply potential scaffolding errors remaining in the dense repeat regions, although interestingly even though the data was more limiting the same pattern appeared to be apparent for the CN115125 genotype, which could suggest a dispersed structure for the active centromeric region^47^. Where comparison was feasible the two genotypes showed a common dichotomy of centromeric regions, with conservation of gene content but rapid divergence in sequence constitution driven by changes in retrotransposon composition.

Recent work in *B. nigra* to uncover centromere specific sequences through their association with CENH3 indicated that unlike its diploid relatives and almost all analysed plant genomes, *B. nigra* contains no tandemly repeated satellite DNA^36^. Similarly, no characteristic tandem repeat was found in the long-read assemblies; however, analyses of assembled full length LTRs revealed recently amplified (< 1 Mya) elements, in particular ALE-LTRs in the Ni100-LR genome (Figure 6). Rapid amplification of the young LTRs in a nested insertion fashion was observed in all of the Ni100 centromeric regions (Supplementary Table 17). Nested TE insertion is a prevalent phenomena among monocots, but has only been identified infrequently among dicots^48^. The detected recent nested insertion events involving a single family suggest that ALE or related LTRs might play an important role in rapid divergence of centromeres in *B. nigra*, similar to that found when comparing the centromeric region of two rice genotypes^49^. Further studies to fully establish the role of these elements in centromere function in *B. nigra* are required and indeed the long read assembly resources developed for the Ni100 genotype could be leveraged as a model for centromere function research in future.

Finally the improved assemblies for all three diploid *Brassica* genomes allowed a median ancestral *Brassica* genome (n =9) to be resolved based on 178 syntenic blocks. The calculated DCJ distance between the genomes reflects the age of divergence between the B genome and A/C genome lineages (Figure 7A). While *B. rapa* and *B. oleracea* have chromosomes which share extensive homology with ancestral chromosomes, the extent of the rearrangements separating the B genome would explain the limited genic exchange that has been possible across the two lineages. Therefore, capturing novel diversity from the third *Brassica* genome for crop improvement strategies in its related species may be more efficient using next generation breeding techniques such as CRISPR/Cas9.

The ability to relatively quickly and affordably generate contiguous genome assemblies provides a platform for developing true pan-genomes for many species. Such assemblies will allow an accurate comparison of not only gene content, but repeat composition and distribution and reveal the range and complexity of structural variation. There are still some limitations; however, with the continuing improvements to the technology and the optimisation of software dedicated to analyses of these new data types, resolution of these problems should be swift.

## Materials and Methods

### Plant material and DNA extraction

*Brassica nigra* CN115125 (C2) and Ni100 were grown in a greenhouse at Agriculture and Agri-Food Canada, Saskatoon Research and Development Centre, under 20/18 °C, 16/8 hour days. Leaf tissue was collected from three week old plants, after two days of dark treatment flash frozen and stored at −70 °C. Nuclear isolation was performed as described in Zhang et al^50^ and high molecular weight DNA was extracted using a modified CTAB method^51^. Briefly, approximately 20 g of leaf tissue was homogenized in 200 mLs ice-cold 1× HBS solution (0.01 M Trizma base, 0.08 M KCL, 1 mM spermidine, 0.01 M EDTA, 0.5 M sucrose, 1 mM spermine plus 0.15% β-mercaptoethanol). Five mLs of 1X HBS plus 20% Triton X-100 were added to the homogenate and mixed slowly with a magnetic stir bar for 1 hour on ice, then filtered through two layers of cheesecloth and one layer of Miracloth. The nuclei were pelted by centrifugation of homogenate at 1800 *g* for 20 min at 4 °C. The pellet was washed by re-suspension in 1 × HB plus 0.5% Triton-×100 on ice and centrifuged at 1800 *g* for 20 min at 4 °C three times to purify the nuclei. The final pelleted nuclei were re-suspended in 10 ml lysis buffer (100 mM TrisHCl, 100 mM NaCl, 50 mM EDTA, 2% CTAB) treated with proteinase K, followed by RNAase A (37°C for 30 min) and high molecular weight DNA was extracted after two cycles of phenol/chloroform clean-up and ethanol precipitation. DNA quality and quantity was measured using an Agilent Bioanalyzer and Qubit fluorometer, respectively.

### Oxford Nanopore sequencing and reads processing

The C2 genome was sequenced on a MinION while the Ni100 genome was sequenced on a GridION. For the C2 genome, 1D (SQK-LSK108) and 1D^2^ (SQK-LSK308) genomic DNA libraries were prepared following the nanopore protocol (https://community.nanoporetech.com/protocols). For size selected DNA, 4 µg of DNA was sheared with a Covaris g-TUBE to obtain 10 kb fragments. Two µg of sheared and un-sheared DNA was used for library preparation for both the 1D and 1D^2^ methods. For the Ni100 genome, 1D (SQK-LSK109) and Rapid (SQK-RAD004) libraries were prepared for sequencing on the GridION. MinION sequencing used MinKnow v1.4.2 with albacore (v1.1.2) live base calling enabled with default parameters. ONT reads with read quality score ≥10 (q10) were filtered from the ONT fastq files (Supplementary Table 3). For the Ni100 genome, sequenced using the GridION, MinKnow 2.0 and live base calling was completed with Guppy and ONT reads with read quality score ≥7 (q7) were used for assembly. Nanostat^52^ was used to compute the sequencing statistics for each run with both raw and quality filtered data.

### Illumina sequencing

Genomic DNA extracted as above was used for whole genome Illumina sequencing. For CN115125, 2 µg of DNA was fragmented using a Covaris sonicator to obtain a 350 bp fragments, and TruSeq DNA PCR-Free library was prepared following the manufacturers protocol (Illumina Inc.). The normalized library was paired-end sequenced in 2×101 bp, and 2×250 bp rapid run mode on HiSeq 2500 platform (Illumina Inc, San Diego, CA, USA. In total, over 82 Gb of short read sequences with ~137x physical coverage were generated for C2 (Supplementary Table 21). For Ni100, whole genome shotgun Illumina paired-end (300-700 bp insert size) and Illumina and Roche/454 (454 Life Sciences, Branford, CT, US) mate-pair libraries (3 - 45 Kb insert size) were developed following the manufacturers protocols. In total 115 Gb (~192x physical coverage) were sequenced and used for whole genome assembly by SOAPdenovo (version 1.05) following the previous approach^16^ (Table1; Supplementary Table 21).

Total RNA was extracted from bud, flower, leaf, seedling, root, and silique tissue samples for Ni100, and from leaf and bud samples for C2 using the RNeasy plant mini kit (QIAGEN, Canada), including on-column DNase digestion (Supplementary Table 21). Total RNA integrity and quantity was assessed on a Bioanalyzer (Agilent). Illumina Tru-Seq RNA-seq libraries were prepared and 125 bp paired-end sequencing was performed using the Illumina Hiseq 2000 platform. A total of 11 Gb and 39 GB raw Illumina RNA-seq data was generated for C2 and Ni100, respectively (Supplementary Table 21). Reads were filtered for low quality (below Q30), adapter sequence, potential PCR duplicates, and length (< 55bp) with Trimmomatic (v.0.32). RSEM^53^ (rsem-calculate-expression) was used to calculate the expression in transcripts per million.

### Genome size estimation based on k-mer analysis

Jellyfish v2.2.6 was used to estimate k-mer frequency distribution based on the subset (~35 Gb) of raw 2×250 PE Illumina reads with a kmer length of 17. Output histogram was uploaded to findGSE to estimate the genome size, heterozygosity and repeat fraction^54^. Analysis has shown that the genome size is about 570 Mb and 607.8 Mb for Ni100 and C2 and was used as a haploid genome size for the study (Supplementary Figure 19).

### Nanopore sequence assembly and polishing

Raw ONT fastq data was filtered for quality at q10 and q7 for C2 and Ni100, respectively, and the resulting reads were error corrected using CANU 1.6 with default parameters^22^. The C2 filtered data was assembled with three different assemblers (SMARTDenovo, wtdbg, Miniasm). Minimap2 was used to generate overlaps of corrected reads with k-value 24 and other default parameters (-csw5 -L100 -m0) followed by assembly using miniasm^22,55^. SMARTDenovo (https://github.com/ruanjue/smartdenovo) was used with the k-value of 24 and recommended parameters. wtdbg tool (https://github.com/ruanjue/wtdbg) was used to assemble the reads with k17 and k24 with default parameters (-H -S 1.02 -e 3). The best assembly for the C2 genome (S4) based on contiguity and genome coverage was selected for further analysis (Supplementary Table 1). Based on this preliminary analyses the Ni100 genome were assembled using SMARTDenovo with kmer 24 and default parameters. Both draft assemblies were polished using eight iterations of PILON^23^ with available Illumina reads.

### Contig Scaffolding

Leaf tissue of C2 was provided to Dovetail genomics (Santa Cruz, CA, USA) who prepared and sequenced CHiCAGO^™^ and Hi-C libraries. The polished assemblies, CHiCAGO^™^, and Dovetail Hi-C library reads were used as input for scaffolding using Dovetail’s HiRise™ pipeline^56^. A modified SNAP read mapper uses CHiCAGO^™^ and HiC reads to align to the draft assembly, HiRise™ produces a likelihood model for the genomic distance between read pairs, computing the optimum threshold to join contigs and to identify putative misjoins.

A genetic map derived from genotyping-by-sequencing data of a backcross population of 72 *B. nigra* lines derived from the cross Ni100/doubled-haploid line A1//Ni100 was used to anchor contigs from all assemblies to the pseudomolecules. 20,689, 19,666 and 21,034 loci were anchored to the genome assemblies of C2, Ni100-SR and Ni100-LR, respectively. The assembly was confirmed using Genome-Ordered Graphical Genotypes (GOGGs)^57^ based on transcriptome re-sequencing of lines from the *B. juncea* VHDH mapping population^58^ and genome re-sequencing of lines from the *B. juncea* YWDH population^59^. The GOGGs also enabled four previously unanchored scaffolds to be incorporated into the chromosome assemblies. Sequences of the RFLP clones used to generate the genetic map in Lagercrantz and Lydiate (1995)^60^ were aligned to the assemblies to name and orient the pseudomolecules accordingly, based on the internationally agreed standard (http://www.Brassica.info). A lookup table comparing chromosome (linkage group) names between the two published nomenclatures for the B genome are shown in supplementary Table 7.

### Assembly quality assessment

Quality of the assembly was estimated using the single-copy orthologous gene analysis (BUSCO v3.0.2)^24^ with Embryophyta OrthoDB version 9. The 1,440 genes were searched in the assembly using Augustus (version 3.2.1)^61^, NCBI’s BLAST (version 2.2.31+)^62^, and HMMER (version 3.1b2) by BUSCO. In addition, genome discrepancies were estimated using qualimap^25^ by mapping the Illumina reads against the polished assembly. Bowtie-2^63^ with default parameters was used for mapping the Illumina reads against the assembly.

### Genome Annotation

RNA-Seq (39 GB) data for Ni100 and C2 were aligned against their respective genome assemblies using STAR v2.7 (max. 3% mis-matches over 95% read length), and subsequently assembled using Trinity (v2.8.4) genome-guided approach with default parameters. In total, 110,767 and 124,851 transcripts were assembled for Ni100 and C2, respectively. The assembled transcripts, along with protein sequences from *A. thaliana*, *A. lyrata*, *B. rapa*, and *B. oleracea*, were used as evidence for the MAKER-P annotation pipeline^64^. Snap and Augustus *ab initio* predictors were configured for use by MAKER-P in hint-based mode utilizing the protein and transcript as input evidence. Approximately 6% of the predicted gene models were found to be mis-joined based on *A. thaliana* gene structure and *B. nigra* transcript evidence, and were split into two or more alternate models. PASA (v2.3.3) software^65^ was then used to further assemble Trinity output and to incorporate the transcript alignment evidence into MAKER gene annotation. In total, 59,852 and 67,021 coding genes were annotated for Ni100 and C2, respectively. Of the annotated genes, 48,621 (81.2%) of Ni100 and 54,586 (81.4%) of C2 gene models have expression values of TPM>0. BLASTP revealed that 55,022 (92.0%) of Ni100 and 59,780 (89.2%) of C2 genes have significant hits (E-value cutoff 10^−5^) against the Uniprot Plant database.

The gene naming convention proposed for *B. rapa* V3^19^ was used with minor modifications: **Bni** (for *Brassica nigra*) followed by the chromosome number with leading zero, and a letter “**g**” (for gene), e.g. **B01g** (for B genome chromosome 1). Six digit gene numbers were assigned in steps of 10 with leading zeros from top to bottom of chromosomes. Following the gene number and separated by a period, to distinguish genome versions and between genotypes, “**2N**” was assigned to Ni100 LR (genome version 2), and “**1C2**” to C2 (genome version 1); for example, BniB01g023500.2N. Low confidence genes were defined as those models with neither transcriptome evidence support nor significant hits to Uniprot Plant database. The low confidence genes were named similarly as described above but with a letter “**p**” to distinguish them.

### Repeat annotation

*A de novo* repeat library was developed using RepeatModeler (Version 1.0.11; http://www.repeatmasker.org/RepeatModeler/) which employ two *de novo* repeat finding programs (RECON and RepeatScout) for identification of repeat families. After removing potential false positives based on the homology with *A. thaliana* gene models, a total of 374 repeat models were retained. In addition, a previously developed repeat library for *B. nigra* Ni100 which contains 950 repetitive elements was merged to develop a final repeat library with 1324 that was used for repeat annotation in the whole genome. *Repeatmasker* was employed to estimate the repeat copies, proportion and distribution into the genome^66666666656463^.

Centromeric location was identified based on the distribution of centromere associated repeats such as Centromere specific retrotransposon of *Brassica* (CRB) and *B. nigra* specific centromere associated repeat (pBN35 - X16588.1)^35,67^. It is expected that that both the CRB and *B. nigra* specific centromere associated repeat were associated with the centromere and adjutant regions. Based on the distribution of these two elements using BLAST, centromere regions were located in the assembly.

Full length long terminal repeat retrotransposons (FL-LTR-RTs) were identified from both genome assemblies using LTR_harvest^68^ and LTR_Finder^69^. The resulting outputs (.scn) were fed into the LTR retriever program^70^ to extract the FL-LTR-RTs. Copy number, distribution and divergence time of the LTR-RT were comparatively analyzed between the two reference genomes. FL-LTR-RTs were classified to different families based on the homology with the repeat library from *Repeat Explorer*^71^.

Phylogenetic analysis was done using the reverse-transcriptase (RT) domain sequences of ALE and OTA elements from the thee *B. nigra* genomes. RT domains obtained from the Pfam database, Accession No. PF07727 and PF00078, were used as a query to search against the FL-LTRs of ALE and OTA sequences, respectively by BLASTx and the best hit with minimum of 200 bp overlap with query sequences were used for further analysis. RT domain sequences of ALE and OTA families from the thee *B. nigra* genomes were aligned separately by the clustalW aligner and a tree was generated using neighbor-joining method with the 500 bootstrap replications by MEGA7. FL-LTR-RTs annotated for the whole genome using the LTR-retriever were manually analyzed to identify nested TE insertion.

### Gene and genome evolution

OrthoFinder v2.2.7^29^ was used to identify members of gene families and assess expansion of gene families in C2 and Ni100-LR, by clustering the annotated genes with closely related species, *B. rapa* ^19^, *B. oleracea*^16^ and *A. thaliana* (Araport 11) (Supplementary Table 10). CAFE v4.2.1^30^ was used to identify expansion, contraction and rapidly evolving gene families among the six genomes based on the orthogroups obtained from the OrthoFinder analysis.

Synteny analysis was performed to identify syntenic genes between *B. nigra* Ni100-LR/C2 and *A. thaliana* using *A. thaliana* proteome (Araport10) as described previously^16^. Briefly, based on best BLASTP values (E-value 1e^−20^ or better) syntenic gene pairs between *B. nigra* and *A. thaliana* were employed in DAGChainer with default parameters to compute the chain score^72^. Manual curation based on better chain score was done to create the final syntelog table (Supplementary Table 21). Tandemly duplicated genes and proximal genes were identified following the previously reported approach^73^. Briefly, potential homologous pairs between each of three genomes were identified by all-versus-all BLASTP with E < 1e-10. Then MCScanX (default parameters) was used to identify duplicated pairs from Ni100 and C2 that formed intra-species syntenic chains. These pairs were set aside and classified as WGD-derived gene pairs. The remaining pairs (or BLASTP hits) are classified as either Tandem (next to each other on the same chromosome) or proximal (>1 and <=10 genes on the same chromosome).

For phylogenetic analysis, a data matrix consisting of 1150 syntelogs retained in *A. thaliana* and each of the sub-genomes (least fractionated: LF; most fractionated 1: MF1; and most fractionated 2: MF2) of *B. rapa, B. oleracea* and *B.nigra* was constructed. Sequences from individual syntelog gene sets were aligned using ClustalW version 2.1; ^74^, and poorly aligned regions were removed using trimAL version 1.2; ^75^. Trimmed sequences were concatenated using the Phyutility program (Smith and Dunn, 2008) to produce the final data matrix comprising a total alignment length of 807,943 bp. The phylogenetic relationship was inferred using maximum likelihood method implemented in RAxML version 8.2.12; ^76^ using rapid bootstrapping (100 replications) and a GTRGAMMA substitution model. The resulting phylogenetic tree was visualized by using the interactive Tree of Life version 4; ^77^ web server.

For ancestral genome reconstruction, given gene orders in a set of genomes G and a distance measure d, the median problem is to find a genome m that minimizes the sum of distances d^Σ^ = Σ_g∈G_ d(m, g). The median problem is known to be NP-hard under the DCJ distance^78^. Given three genomes and di,j as the pairwise DCJ distance between genome i and genome j, the metric dΣ has the following property, the lower bound is 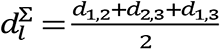, and the upper bound is 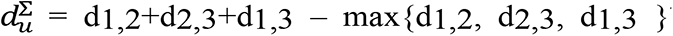^41^. Therefore, given the pairwise DCJ distances calculated for the three genomes, *B. rapa*, *B. nigra* and *B. oleracea*, 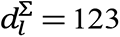 and 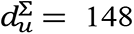. The ASMedian-Linear algorithm^41^ is designed to find exact solutions to the DCJ median problem on multi-chromosomal genomes. It uses a divide-and-conquer approach to decompose the multiple breakpoint graph to its “adequate” subgraphs, find optimal solutions for its parts, then combine the optimal solutions. A total of 25,866 orthologous genes were identified between the genomes of *B. rapa*, *B. nigra* and *B. oleracea*, These genes were used to identify 178 unique blocks where although distance between blocks was estimated based on fractionated genes, block reversals were used to fix block breakpoints. Finally, a median genome was calculated using the ASMedian-Linear algorithm as nine ancestral chromosomes with a genome size of 321 Mb.

Ks analysis (distribution of synonymous substitutions) was performed as described previously^12^. Briefly, for each pair of syntelogs between the *Brassica* species or the subgenomes of each *Brassica* species, protein sequences were aligned using ClustalW version 2.1; ^74^ and the corresponding codon alignments were produced using PAL2NAL ^79^. Ks values for each sequence pair were calculated using the maximum likelihood method implemented in codeml of the PAML package^80^ under the F3×4 model ^81^. Histograms were generated using the log transformed Ks values >0.001. Gaussian mixture models were fitted to the ln (Ks) values using the R package Mclust, and the number of Gaussian components, the mean of each component and fractions of data were calculated. The Bayesian Information Criterion was used to determine the best fitting model to the data. The fit of the determined models were confirmed by χ2 tests.

The presence of resistance (R) genes were identified using the RGAugury pipeline (version 2017.10.21)^82^; transcription factors (TFs) and transcription regulators (TRs) and protein-kinase families were identified by iTAK (current version: v1.7 - 05/13/16)^83^.

### Structural variants analysis

SVs such as insertions, deletions, inversions, duplications, and translocations were identified using both Ni100-LR and C2-LR assemblies. Raw long reads of both genomes were mapped using NGMLR long-read aligner on Ni100-LR as a reference and SVs were called using Sniffles with minimum read depth of 20^31^. Likewise, SVs are predicted using C2-LR as a reference assembly. Furthermore, cross-validation of SVs identified by Sniffles was done using another SV identifier, Picky, with the same read depth of 20^32^. SVs shared by both callers were identified as high-quality SVs and used for further analysis.

### Whole genome Bisulfite sequencing (WGBS)

Genomic DNA was isolated from leaf tissue of *B. nigra* Ni100 with two biological replications and leaf and bud tissue of *B. nigra* CN115125 nuclei using Qiagen’s DNeasy Plant Kit (Qiagen GmbH, Hilden, Germany) following the manufacturer’s protocol. Zymo Research EZ DNA Methylation kit was used for bisulfite conversion on 100 ng of DNA along with 0.5% w/w of unmethylated lambda DNA (Promega, US), included to evaluate bisulfite conversion efficiency. Library construction was performed according to the Illumina TruSeq® DNA Methylation Kit Reference Guide (15066014 v01). The libraries were quantified as above and PE sequenced (2 x 125 bp) using an Illumina HiSeq 2000.

Quality filtered WGBS reads were used to analyze cytosine methylation ratios following alignment using BSMAP (v2.9) (Supplementary Table 21)^84^. Lambda DNA was included in each library as a control to estimate bisulfite conversion efficiency. In all instances the conversion rate was estimated to exceed 99%. The evidence to assign the methylatation status of each cytosine surveyed was determined by using the binomial probability distribution. Methylation patterns were determined and summarized using the support from available genome annotation. Methylation patterns were partitioned by context (CG, CHH, CHG) reflecting the underlying biochemistry underpinning their maintenance. Statistical relationships and data organization was performed using custom Perl and R scripts with the support from Datatable, dplyr, stringr, genomation and MethylKit, All graphical summaries were developed using ggplot2.

### CpG context in nanopore reads by Nanopolish

Since, nanopore reads have an ability to output signals for methylated and unmethylated cytosine bases, Nanopolish was used to detect the CpG context in the whole genome of Ni100^33^. Nanopolish 0.10.1 was used to call bases for methylated and unmethylated from raw nanopore reads and results were filtered as described based on either log-likelihood ratio or read depth.

### Data availability

Genome assembly and annotation, DNA-seq data, RNA-seq data and WGBS-seq data has deposited to NCBI under BioProject ID PRJNA436722. A Jbrowse instance for each genome can be accessed at http://cruciferseq.ca.

## Acknowledgements

This research was supported by funding from the Agriculture and Agri-Food Canada Canadian Crop Genomics Initiative and through the Plant Phenotyping and Imaging Research Centre (P^2^IRC) funded by the Canada First Research Excellence Fund and managed by the Global Institute for Food Security. Sampath Perumal was additionally supported by a MITACS elevate post-doctoral fellowship.

## Contributions

S.P., A.G.S and I.A.P.P conceived the study. S.P. and E.H. performed ONT sequencing. L.T and Z.K developed the backcross mapping population and linkage map. Z.H and I.B carried out additional genetic confirmation of scaffold/contig order. C.K., S.P., L.J, M.B, and I.A.P.P carried out assembly, bioinformatic and statistical analyses. S.K performed Ks-based divergence analysis. L.J, C.Z, and D.S performed ancestral genome characterization. K.N.H and S.J.R performed bisulfite sequencing and methylome analysis. B.C provided additional RNASeq data. S.P and I.A.P.P wrote the manuscript. All authors read and contributed to the final manuscript.

## Supplementary information

Supplementary Tables 1-21 (Excel file).

Supplementary Figures 1-19 (.pdf).

## References

1. Bevan MW, Uauy C, Wulff BB, Zhou J, Krasileva K, Clark MD. Genomic innovation for crop improvement. Nature 543, 346 (2017).

2. Abberton M, et al. Global agricultural intensification during climate change: a role for genomics. Plant biotechnology journal 14, 1095–1098 (2016).

3. Scheben A, Wolter F, Batley J, Puchta H, Edwards D. Towards CRISPR/Cas crops–bringing together genomics and genome editing. New Phytologist 216, 682–698 (2017).

4. Michael TP. Plant genome size variation: bloating and purging DNA. Briefings in functional genomics 13, 308–317 (2014).

5. Lim KB, et al. Characterization of the centromere and peri1centromere retrotransposons in Brassica rapa and their distribution in related Brassica species. The Plant Journal 49, 173–183 (2007).

6. Sedlazeck FJ, Lee H, Darby CA, Schatz MC. Piercing the dark matter: bioinformatics of long-range sequencing and mapping. Nature Reviews Genetics 19, 329 (2018).

7. Jiao W-B, Schneeberger K. The impact of third generation genomic technologies on plant genome assembly. Current opinion in plant biology 36, 64–70 (2017).

8. Sedlazeck FJ, Lee H, Darby CA, Schatz MC. Piercing the dark matter: bioinformatics of long-range sequencing and mapping. Nature Reviews Genetics 19, 329–346 (2018).

9. Koren S, Phillippy AM. One chromosome, one contig: complete microbial genomes from long-read sequencing and assembly. Current opinion in microbiology 23, 110–120 (2015).

10. Jiao Y, et al. Improved maize reference genome with single-molecule technologies. Nature 546, 524 (2017).

11. Deamer D, Akeson M, Branton D. Three decades of nanopore sequencing. Nature Biotechnology 34, 518 (2016).

12. Kagale S, et al. Polyploid evolution of the Brassicaceae during the Cenozoic era. The Plant cell 26, 2777–2791 (2014).

13. Lysak MA, Koch MA, Pecinka A, Schubert I. Chromosome triplication found across the tribe Brassiceae. Genome research 15, 516–525 (2005).

14. U N. Genome analysis in Brassica with special reference to the experimental formation of B. napus and peculiar mode of fertilization. Jap J Bot 7, 389–452 (1935).

15. Truco MJ, Quiros CF. Structure and organization of the B genome based on a linkage map in Brassica nigra. Theoretical and Applied Genetics 89, 590–598 (1994).

16. Parkin IA, et al. Transcriptome and methylome profiling reveals relics of genome dominance in the mesopolyploid Brassica oleracea. Genome biology 15, R77 (2014).

17. Liu S, et al. The Brassica oleracea genome reveals the asymmetrical evolution of polyploid genomes. Nature communications 5, 3930 (2014).

18. Chalhoub B, et al. Early allopolyploid evolution in the post-Neolithic Brassica napus oilseed genome. Science 345, 950–953 (2014).

19. Zhang L, et al. Improved Brassica rapa reference genome by single-molecule sequencing and chromosome conformation capture technologies. Horticulture Research 5, 50 (2018).

20. Yang J, et al. The genome sequence of allopolyploid Brassica juncea and analysis of differential homoeolog gene expression influencing selection. Nature genetics 48, 1225 (2016).

21. Belser C, et al. Chromosome-scale assemblies of plant genomes using nanopore long reads and optical maps. Nature Plants 4, 879–887 (2018).

22. Koren S, Walenz BP, Berlin K, Miller JR, Bergman NH, Phillippy AM. Canu: scalable and accurate long-read assembly via adaptive k-mer weighting and repeat separation. Genome research, gr. 215087.215116 (2017).

23. Walker BJ, et al. Pilon: An Integrated Tool for Comprehensive Microbial Variant Detection and Genome Assembly Improvement. PloS one 9, e112963 (2014).

24. Simão FA, Waterhouse RM, Ioannidis P, Kriventseva EV, Zdobnov EM. BUSCO: assessing genome assembly and annotation completeness with single-copy orthologs. Bioinformatics 31, 3210–3212 (2015).

25. Okonechnikov K, Conesa A, García-Alcalde F. Qualimap 2: advanced multi-sample quality control for high-throughput sequencing data. Bioinformatics 32, 292–294 (2015).

26. Golicz AA, et al. The pangenome of an agronomically important crop plant Brassica oleracea. Nature communications 7, 13390 (2016).

27. Wu TD, Watanabe CKJB. GMAP: a genomic mapping and alignment program for mRNA and EST sequences. 21, 1859–1875 (2005).

28. Bachmann JA, Tedder A, Laenen B, Steige KA, Slotte TJGG, Genomes, Genetics. Targeted long-read sequencing of a locus under long-term balancing selection in Capsella. 8, 1327–1333 (2018).

29. Emms DM, Kelly SJGb. OrthoFinder: solving fundamental biases in whole genome comparisons dramatically improves orthogroup inference accuracy. 16, 157 (2015).

30. Han MV, Thomas GW, Lugo-Martinez J, Hahn MWJMb, evolution. Estimating gene gain and loss rates in the presence of error in genome assembly and annotation using CAFE 3. 30, 1987–1997 (2013).

31. Sedlazeck FJ, et al. Accurate detection of complex structural variations using single-molecule sequencing. Nature methods 15, 461–468 (2018).

32. Gong L, et al. Picky comprehensively detects high-resolution structural variants in nanopore long reads. Nature methods 15, 455–460 (2018).

33. Simpson JT, Workman RE, Zuzarte PC, David M, Dursi LJ, Timp W. Detecting DNA cytosine methylation using nanopore sequencing. Nature methods 14, 407 (2017).

34. Koo DH, et al. Rapid divergence of repetitive DNAs in Brassica relatives. Genomics 97, 173–185 (2011).

35. Lim KB, et al. Characterization of the centromere and peri-centromere retrotransposons in Brassica rapa and their distribution in related Brassica species. The Plant journal : for cell and molecular biology 49, 173–183 (2007).

36. Wang G-x, et al. ChIP-cloning analysis uncovers centromere-specific retrotransposons in Brassica nigra and reveals their rapid diversification in Brassica allotetraploids. Chromosoma 128, 119–131 (2019).

37. Lysak MA, Mandáková T, Schranz MEJCoipb. Comparative paleogenomics of crucifers: ancestral genomic blocks revisited. 30, 108–115 (2016).

38. Cheng F, et al. Biased Gene Fractionation and Dominant Gene Expression among the Subgenomes of Brassica rapa. PloS one 7, e36442 (2012).

39. Eichler EE, Sankoff DJs. Structural dynamics of eukaryotic chromosome evolution. 301, 793–797 (2003).

40. Yancopoulos S, Attie O, Friedberg RJB. Efficient sorting of genomic permutations by translocation, inversion and block interchange. 21, 3340–3346 (2005).

41. Xu AW. DCJ median problems on linear multichromosomal genomes: graph representation and fast exact solutions. In: RECOMB International Workshop on Comparative Genomics (ed^(eds). Springer (2009).

42. Michael TP, et al. High contiguity Arabidopsis thaliana genome assembly with a single nanopore flow cell. Nature communications 9, 541 (2018).

43. Gabur I, Chawla HS, Snowdon RJ, Parkin IAP. Connecting genome structural variation with complex traits in crop plants. Theoretical and Applied Genetics, (2018).

44. Cameron DL, Di Stefano L, Papenfuss AT. Comprehensive evaluation and characterisation of short read general-purpose structural variant calling software. Nature communications 10, 3240 (2019).

45. De Coster W, et al. Structural variants identified by Oxford Nanopore PromethION sequencing of the human genome. Genome research, (2019).

46. van Dijk EL, Jaszczyszyn Y, Naquin D, Thermes C. The third revolution in sequencing technology. Trends in Genetics 34, 666–681 (2018).

47. Muller H, Gil Jr J, Drinnenberg IAJTiG. The impact of centromeres on spatial genome architecture. (2019).

48. Kronmiller BA, Wise RPJPp. Computational finishing of large sequence contigs reveals interspersed nested repeats and gene islands in the rf1-associated region of maize. 151, 483–495 (2009).

49. Gao D, Jiang N, Wing RA, Jiang J, Jackson SA. Transposons play an important role in the evolution and diversification of centromeres among closely related species. Frontiers in plant science 6, (2015).

50. Zhang HB, Zhao X, Ding X, Paterson AH, Wing RA. Preparation of megabase1size DNA from plant nuclei. The Plant Journal 7, 175–184 (1995).

51. Allen G, Flores-Vergara M, Krasynanski S, Kumar S, Thompson WJNp. A modified protocol for rapid DNA isolation from plant tissues using cetyltrimethylammonium bromide. 1, 2320 (2006).

52. De Coster W, D’Hert S, Schultz DT, Cruts M, Van Broeckhoven C. NanoPack: visualizing and processing long-read sequencing data. Bioinformatics 34, 2666–2669 (2018).

53. Li B, Dewey CNJBb. RSEM: accurate transcript quantification from RNA-Seq data with or without a reference genome. 12, 323 (2011).

54. Sun H, Ding J, Piednoël M, Schneeberger K. findGSE: estimating genome size variation within human and Arabidopsis using k-mer frequencies. Bioinformatics 34, 550–557 (2018).

55. Li H. Minimap and miniasm: fast mapping and de novo assembly for noisy long sequences. Bioinformatics 32, 2103–2110 (2016).

56. Moll KM, et al. Strategies for optimizing BioNano and Dovetail explored through a second reference quality assembly for the legume model, Medicago truncatula. BMC genomics 18, 578 (2017).

57. He Z, Bancroft IJNg. Organization of the genome sequence of the polyploid crop species Brassica juncea. 50, 1496–1497 (2018).

58. Ramchiary N, et al. Mapping of yield influencing QTL in Brassica juncea: implications for breeding of a major oilseed crop of dryland areas. 115, 807–817 (2007).

59. Guo S, et al. A genetic linkage map of Brassica carinata constructed with a doubled haploid population. 125, 1113–1124 (2012).

60. Lagercrantz U, Lydiate DJJG. RFLP mapping in Brassica nigra indicates differing recombination rates in male and female meioses. 38, 255–264 (1995).

61. Stanke M, Waack S. Gene prediction with a hidden Markov model and a new intron submodel. Bioinformatics 19, ii215–ii225 (2003).

62. Camacho C, et al. BLAST+: architecture and applications. BMC bioinformatics 10, 421 (2009).

63. Langmead B, Salzberg SL. Fast gapped-read alignment with Bowtie 2. Nature methods 9, 357–359 (2012).

64. Campbell MS, Holt C, Moore B, Yandell M. Genome annotation and curation using MAKER and MAKER1P. Current Protocols in Bioinformatics 48, 4.11. 11–14.11. 39 (2014).

65. Haas BJ, et al. Improving the Arabidopsis genome annotation using maximal transcript alignment assemblies. 31, 5654–5666 (2003).

66. Tarailo□Graovac M, Chen N. Using RepeatMasker to identify repetitive elements in genomic sequences. Current Protocols in Bioinformatics, 4.10. 11–14.10. 14 (2009).

67. Schelfhout CJ, Snowdon R, Cowling WA, Wroth JM. A PCR based B-genome-specific marker in Brassica species. Theoretical and Applied Genetics 109, 917–921 (2004).

68. Ellinghaus D, Kurtz S, Willhoeft U. LTRharvest, an efficient and flexible software for de novo detection of LTR retrotransposons. BMC bioinformatics 9, 18 (2008).

69. Xu Z, Wang H. LTR_FINDER: an efficient tool for the prediction of full-length LTR retrotransposons. Nucleic acids research 35, W265–268 (2007).

70. Ou S, Jiang N. LTR_retriever: A Highly Accurate and Sensitive Program for Identification of Long Terminal Repeat Retrotransposons. Plant physiology 176, 1410–1422 (2018).

71. Neumann P, Novák P, Hoštáková N, Macas J. Systematic survey of plant LTR-retrotransposons elucidates phylogenetic relationships of their polyprotein domains and provides a reference for element classification. Mobile DNA 10, 1 (2019).

72. Haas BJ, Delcher AL, Wortman JR, Salzberg SL. DAGchainer: a tool for mining segmental genome duplications and synteny. Bioinformatics 20, 3643–3646 (2004).

73. Qiao X, et al. Gene duplication and evolution in recurring polyploidization–diploidization cycles in plants. Genome biology 20, 38 (2019).

74. Larkin MA, et al. Clustal W and Clustal X version 2.0. Bioinformatics 23, 2947–2948 (2007).

75. Capella-Gutierrez S, Silla-Martinez JM, Gabaldon T. trimAl: a tool for automated alignment trimming in large-scale phylogenetic analyses. Bioinformatics 25, 1972–1973 (2009).

76. Stamatakis A. RAxML version 8: a tool for phylogenetic analysis and post-analysis of large phylogenies. Bioinformatics 30, 1312–1313 (2014).

77. Letunic I, Bork P. Interactive Tree Of Life (iTOL) v4: recent updates and new developments. Nucleic acids research 47, W256–W259 (2019).

78. Tannier E, Zheng C, Sankoff DJBb. Multichromosomal median and halving problems under different genomic distances. 10, 120 (2009).

79. Suyama M, Torrents D, Bork P. PAL2NAL: robust conversion of protein sequence alignments into the corresponding codon alignments. Nucleic acids research 34, W609–612 (2006).

80. Yang Z. PAML 4: phylogenetic analysis by maximum likelihood. Mol Biol Evol 24, 1586–1591 (2007).

81. Goldman N, Yang Z. A codon-based model of nucleotide substitution for protein-coding DNA sequences. Mol Biol Evol 11, 725–736 (1994).

82. Li P, Quan X, Jia G, Xiao J, Cloutier S, You FM. RGAugury: a pipeline for genome-wide prediction of resistance gene analogs (RGAs) in plants. BMC genomics 17, 852 (2016).

83. Zheng Y, et al. iTAK: A Program for Genome-wide Prediction and Classification of Plant Transcription Factors, Transcriptional Regulators, and Protein Kinases. Molecular plant 9, 1667–1670 (2016).

84. Xi Y, Li W. BSMAP: whole genome bisulfite sequence MAPping program. BMC bioinformatics 10, 232 (2009).

